# Pair-complete structural gating enables Pareto optimization of *de novo* cyclic peptides with preserved binding geometry and improved developability

**DOI:** 10.64898/2026.09.24.754053

**Authors:** Yuanchao Hou, Yixin Zhang, Zhuoyan Liu, Xinfa Peng, Ziye Liu, Miyesier Yusupujiang, You Xu, Jiaxin Hu, Zhenming Liu, Shengwen Yang

**Affiliations:** Key Laboratory of Xinjiang Endemic Phytomedicine Resources, Ministry of Education; School of Pharmacy, Shihezi University, Shihezi 832002, China; The University of Melbourne, Victoria 3010, Australia; Beijing Advanced Center of Cellular Homeostasis and Aging-Related Diseases, Institute of Advanced Clinical Medicine, Peking University, Beijing 100191, China; State Key Laboratory of Natural and Biomimetic Drugs, School of Pharmaceutical Sciences, Peking University, Beijing 100191, China

## Abstract

Cyclic peptides can engage extended protein interfaces, but sequence changes that improve developability often disrupt the selected target-bound geometry. Existing property-guided design methods lack a principled way to preserve this geometry during local multiproperty optimization. Here we present RF-PepTune, which separates structural eligibility from property prioritization. Before any property score is queried, RF-PepTune evaluates single substitutions and all position-distinct pairs induced by admitted substitutions to build a parent-specific, pair-complete admission graph. Pareto Monte Carlo tree search then optimizes predicted permeability, solubility, non-fouling, and low-haemolysis scores exclusively within this graph, and selected sequences undergo full structural re-evaluation. In a matched three-arm comparison of 334 parents across MDM2, GABARAP, and MCL1, joint property–structure success rose from 10.2% without admission to 51.5% with single-substitution admission and 77.5% with pair-complete admission; pair-complete gating improved success by 26.0 percentage points over single-substitution gating (95% CI, 20.7–31.4). Property-positive but structure-failing endpoints fell from 88.3% to 9.3%. In a separate target-balanced set, all 90 prioritized outputs improved all four model-native scores while satisfying prespecified structural criteria. This work establishes pairwise structural gating as a general strategy for constrained multi-objective molecular optimization.

## Introduction

Protein–protein interfaces are frequently large, shallow, and devoid of the deep, well-defined pockets that conventional small molecules exploit. Cyclic peptides occupy a distinctive chemical space between small molecules and biologics: their extended interaction surfaces can engage flat protein interfaces, their conformational restriction can enhance proteolytic stability, and certain macrocyclic scaffolds can cross membranes passively ^1–3^. These attributes make cyclic peptides attractive modalities for targets that have been difficult to drug. Yet obtaining a cyclic peptide with a plausible target-bound conformation does not complete the design problem. Structurally credible binders may still suffer from poor solubility or permeability, non-specific interactions, or hemolytic liability. More critically, sequence changes introduced to improve these developability properties can disrupt the very binding geometry that was initially selected.

Recent deep-learning methods have greatly expanded the accessible design space for target-bound cyclic peptides. Protein-backbone diffusion and inverse-folding models provide complementary capabilities for de novo backbone generation and sequence design ^4–6^. Building on these components, AfCycDesign supports structure prediction, sequence redesign, and de novo design of cyclic peptides, while RFpeptides combines target-conditioned macrocycle generation, sequence design, and structural filtering to produce experimentally validated protein-binding macrocycles ^7,8^. Diffusion, reinforcement-learning, and all-atom generative frameworks have further broadened target-conditioned cyclic-peptide design ^9–13^. In parallel, property-guided approaches optimize membrane permeability and other developability-related objectives ^14–18^, and several methods incorporate structure prediction or structural feedback directly into sequence generation or search ^17,19^. Together, these advances provide powerful foundations for de novo binder generation and property optimization. However, they leave open a central question: how can a selected target-bound geometry be preserved during local multiproperty refinement?

This question is particularly challenging because structural compatibility is not additive. Two substitutions can each satisfy structural criteria when introduced separately, yet fail those criteria when combined. The compatibility of mutation combinations is therefore a distinct consideration in local sequence refinement, not a simple consequence of single-substitution behavior. A related question follows: can pairwise structural evidence be used to define which higher-order sequence states should enter a subsequent property search? Separating this eligibility decision from property-based prioritization provides a principled way to evaluate the contribution of structural admission under matched search conditions.

Here we present RF-PepTune, which integrates RFpeptides-based backbone generation and LigandMPNN sequence design with structure-compatible multiproperty refinement (Fig. 1). Single-substitution screening and subsequent evaluation of position-distinct admitted substitution pairs define a parent-specific, pair-complete admission graph. Pareto Monte Carlo tree search (Pareto-MCTS) then explores this graph using parent-relative PepTune scores for permeability, solubility, non-fouling, and low hemolysis. Selected full sequences undergo structural re-evaluation. We assessed structural admission in a matched three-arm comparison of 334 parents across MDM2, GABARAP, and MCL1. The comparison used common search and endpoint-selection settings to evaluate how single-substitution and pair-complete admission affect joint model-defined property–structure success. By decoupling structural eligibility from property ranking, RF-PepTune identifies candidates with improved predicted property scores and model-defined compatibility with the selected binding geometry, establishing pairwise structural gating as a general strategy for constrained multi-objective optimization of target-bound peptides.

**Fig. 1.**
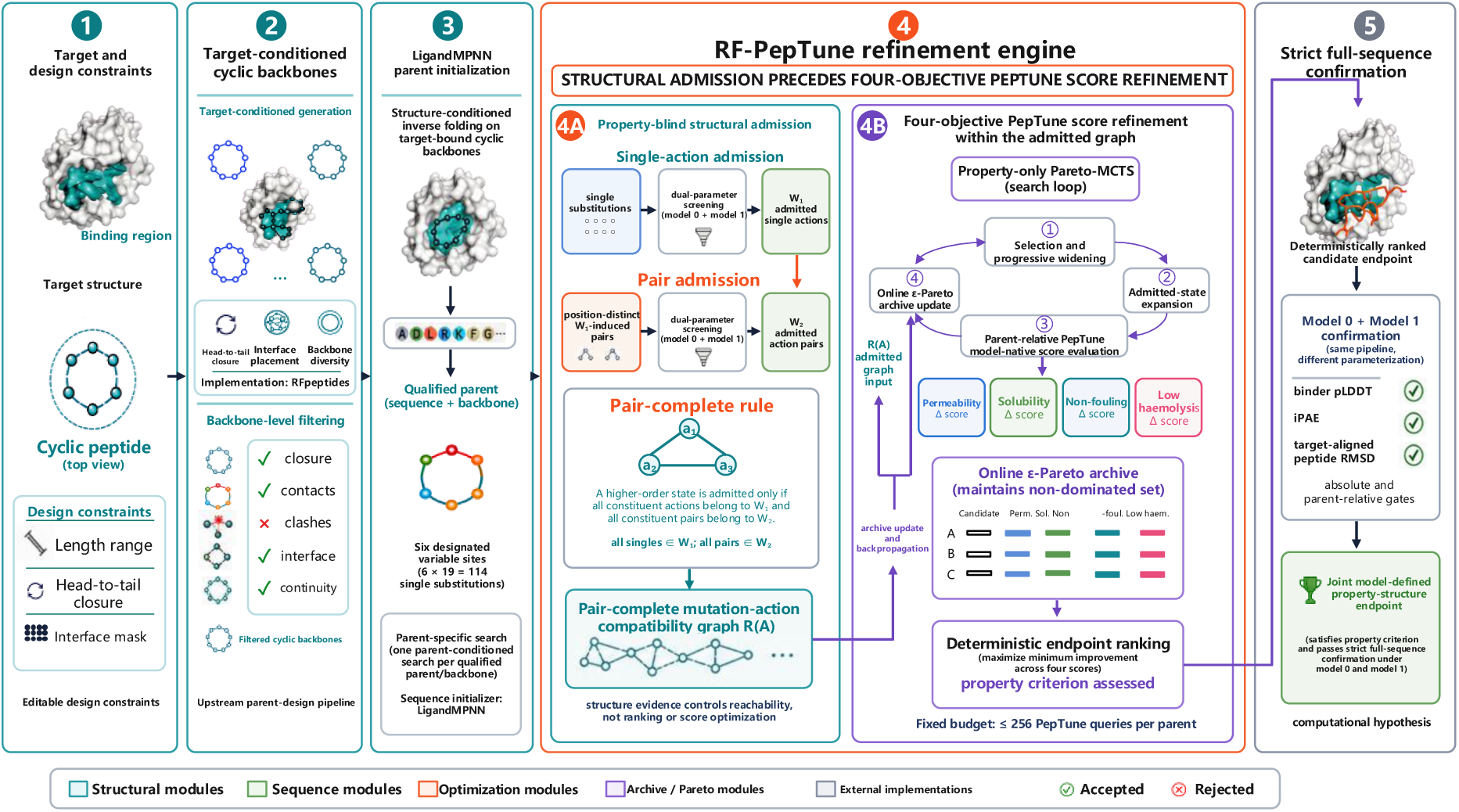
Overview of RF-PepTune. Target-conditioned cyclic backbones are generated and structurally filtered, and LigandMPNN assigns one parent sequence to each qualified backbone. For each parent, six mutable positions define 114 canonical non-parent single substitutions. All 114 substitutions are screened under two parameterizations of the structural workflow (model 0 and model 1) to define the single-substitution admission set W₁. All position-distinct pairs induced by W₁ are then screened under the same structural gates to define W₂. A higher-order state is admitted only when every constituent substitution belongs to W₁ and every constituent pair belongs to W₂, yielding a parent-specific, pair-complete compatibility graph ℛ(A). Pareto Monte Carlo tree search (Pareto-MCTS) then searches only this admitted graph using parent-relative PepTune scores for permeability, solubility, non-fouling, and low hemolysis, retaining non-dominated states in an online ε-Pareto archive. A prespecified max–min rule selects one endpoint, which undergoes full-sequence structural assessment under both parameterizations. Model 0 and model 1 denote two parameterizations of the same ColabDesign/AlphaFold-Multimer workflow and provide a stringent within-workflow criterion, not independent structural validation. Blue, structural screening; green, property-guided search; orange, full-sequence structural assessment.

## Results

### RF-PepTune separates structural eligibility from property ranking to define a pair-complete search space

RF-PepTune treats structural eligibility and property prioritization as two separate decisions (Fig. 1). Target-conditioned cyclic backbones are first generated and filtered, after which LigandMPNN assigns one parent sequence to each qualified backbone. For each parent, six mutable positions define 114 canonical non-parent single substitutions. These substitutions are enumerated before any property score is computed, so that the subsequent search space is defined solely by structural evidence.

The first structural stage evaluates all 114 single substitutions under two parameterizations of the same structural workflow (model 0 and model 1) to define the single-substitution admission set W₁. The second stage evaluates every position-distinct pair induced by W₁ under the same structural gates to define W₂. A higher-order state is admitted only when all its constituent substitutions belong to W₁ and all its constituent pairs belong to W₂. This rule yields a parent-specific, pair-complete compatibility graph ℛ(A), in which higher-order combinations are supported by both singleton and pairwise structural evidence. Across the 334 parents analyzed here, this enumeration produced 204,186 distinct W₁-induced candidate pairs and 408,372 candidate-pair-by-parameterization evaluation records (Supplementary Table 1). Because W₁ and W₂ are constructed before property scoring, structural eligibility is fixed independently of the four developability objectives.

Pareto Monte Carlo tree search (Pareto-MCTS) then searches exclusively within ℛ(A), with at most 256 PepTune queries per parent. States are ranked by parent-relative changes in the four PepTune model-native scores—permeability, solubility, non-fouling, and low hemolysis—and retained in an online ε-Pareto archive. A prespecified max–min rule selects one endpoint for full-sequence structural assessment under both parameterizations. This architecture ensures that structural evidence only defines which states are reachable, whereas property models only prioritize states within that reachable set. The separation makes it possible to evaluate the contribution of structural admission under matched search and endpoint-selection conditions, as described below.

### Pair-complete admission increases joint property–structure success by 26 percentage points

In the matched three-arm experiment, joint model-defined property–structure success—defined as positive parent-relative changes in all four PepTune scores together with passing full-sequence structural assessment under both parameterizations—was achieved for 259 of 334 parents (77.5%) under pair-complete admission, 172 of 334 (51.5%) under W₁-only admission, and 34 of 334 (10.2%) without structural admission (Fig. 2a and Table 1). Relative to W₁-only admission, pair-complete admission increased joint success by 26.0 percentage points (paired-bootstrap 95% CI, 20.7–31.4; Holm-adjusted exact McNemar *P* = 1.33 × 10⁻¹⁸). The corresponding contrast with ungated search was 67.4 percentage points (95% CI, 62.3–72.5; Holm-adjusted exact McNemar *P* = 7.42 × 10⁻⁶⁸).

**Fig. 2.**
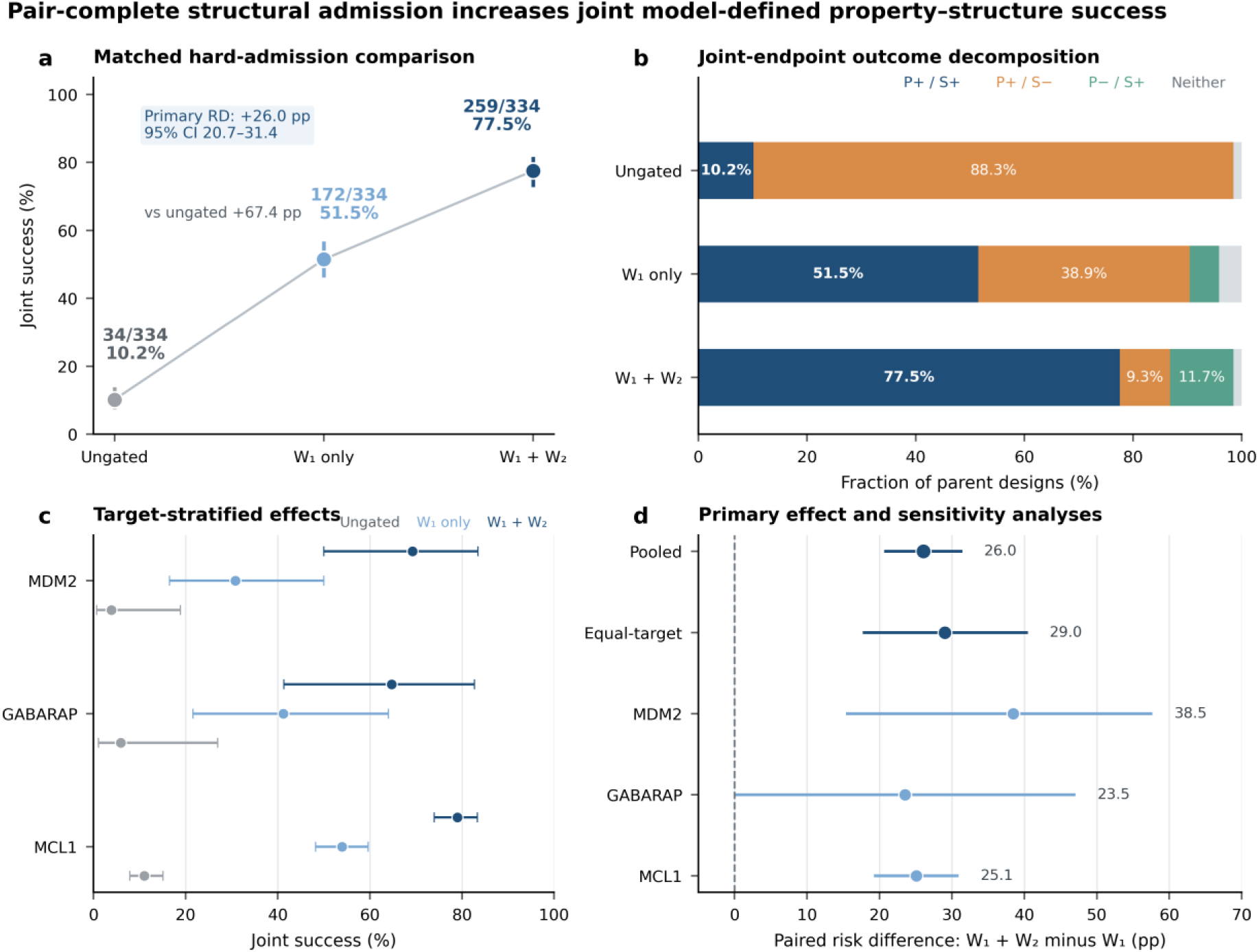
Pair-complete structural admission increases joint model-defined property–structure success under matched search conditions. **a**, Primary-endpoint (joint property–structure success) rates for the three admission arms across 334 parent/source-backbone identities per arm. Ungated, no structural admission; W₁-only, single-substitution admission; W₁+W₂, pair-complete admission. Error bars are two-sided 95% Wilson intervals. b, Four mutually exclusive endpoint categories per arm, defined by positive parent-relative changes in all four PepTune model-native scores (permeability, solubility, non-fouling, and low hemolysis; property +/−) and by prespecified full-sequence structural assessment under both structural parameterizations (model 0 and model 1; structure +/−). Property+ requires positive changes in all four scores; structure+ requires passing all structural gates under both parameterizations. Categories sum to 100% within each arm. **c,** Joint-success rates stratified by target (MDM2, GABARAP, MCL1). **d,** Paired risk differences for W_1_+W_2_ versus W_1_-only admission; positive values indicate higher joint success under pair-complete admission. Points are point estimates and intervals are 95% percentile confidence intervals from 100,000 parent-level bootstrap replicates preserving paired arm outcomes. The primary endpoint required positive changes in all four PepTune scores and passing full-sequence structural assessment under both parameterizations. Parents with no reachable candidate endpoint remained in the prespecified denominator and were counted as failures. n = 334 parents per arm. PepTune scores are computational predictions, not experimental measurements.

**Table 1.** Target-specific and pooled joint model-defined property–structure success in the matched hard-admission comparison.

| Target | n | Ungated n (%) | W<sub>1</sub> only n (%) | W<sub>1</sub>+W<sub>2</sub> n (%) | Paired RD vs W<sub>1</sub>, pp (95% CI) |
| --- | --- | --- | --- | --- | --- |
| MDM2 | 26 | 1 (3.8%) | 8 (30.8%) | 18 (69.2%) | +38.5 (15.4–57.7) |
| GABARAP | 17 | 1 (5.9%) | 7 (41.2%) | 11 (64.7%) | +23.5 (0.0–47.1) |
| MCL1 | 291 | 32 (11.0%) | 157 (54.0%) | 230 (79.0%) | +25.1 (19.2–30.9) |
| Pooled | 334 | 34 (10.2%) | 172 (51.5%) | 259 (77.5%) | +26.0 (20.7–31.4) |

RD, paired risk difference in percentage points. Confidence intervals are 100,000-replicate paired-bootstrap percentile intervals. The pooled row is the prespecified 334-parent analysis; target-specific rows are supportive.

Endpoint composition differed across the three admission policies (Fig. 2b). Among the 334 parents, the selected endpoint was property-positive but structure-failing for 88.3% under ungated search, 38.9% under W₁-only admission, and 9.3% under pair-complete admission. The overall property-positive endpoint rate was 98.5% under ungated search and 86.8% under pair-complete admission. Thus, pair-complete admission reduced both the frequency of property-positive structural failures and the overall rate of property-positive endpoints. These frequencies summarize the endpoints selected under the matched property-query budget.

Paired risk-difference estimates were positive for MDM2 (+38.5 percentage points), GABARAP (+23.5 percentage points), and MCL1 (+25.1 percentage points), and the equal-target estimate was +29.0 percentage points (95% CI, 17.7–40.5; Fig. 2c,d). The equal-target estimate differs slightly from the pooled estimate because MCL1 contributed 291 of the 334 parents, whereas MDM2 and GABARAP contributed 26 and 17, respectively. The three arms shared the same parents, mutable positions, PepTune scorer, Pareto-MCTS implementation, query budget, parent-specific random seeds, endpoint-ranking rule, and full-sequence structural-assessment protocol. Hard structural admission was therefore the only arm-level experimental variable.

Across six post hoc structural-clustering definitions—combining peptide Cα RMSD thresholds of 0.5, 1.0, or 2.0 Å with or without a contact-fingerprint criterion and stratified by target and peptide length—all stratified cluster-bootstrap 95% CIs for the W₁+W₂ versus W₁-only risk difference remained above zero (Supplementary Table 3). These analyses supported the stability of the positive contrast under the examined clustering definitions.

### Pairwise screening excludes incompatible substitution pairs and contracts higher-order reachability

Pairwise structural screening provided information beyond the constituent singleton decisions. Among position-distinct pairs induced by W₁, W₂ excluded median fractions of 29.3% for MDM2, 34.5% for GABARAP, and 25.1% for MCL1 (Fig. 3a). Thus, substitutions that were individually admitted did not necessarily form an admissible pair.

**Fig. 3.**
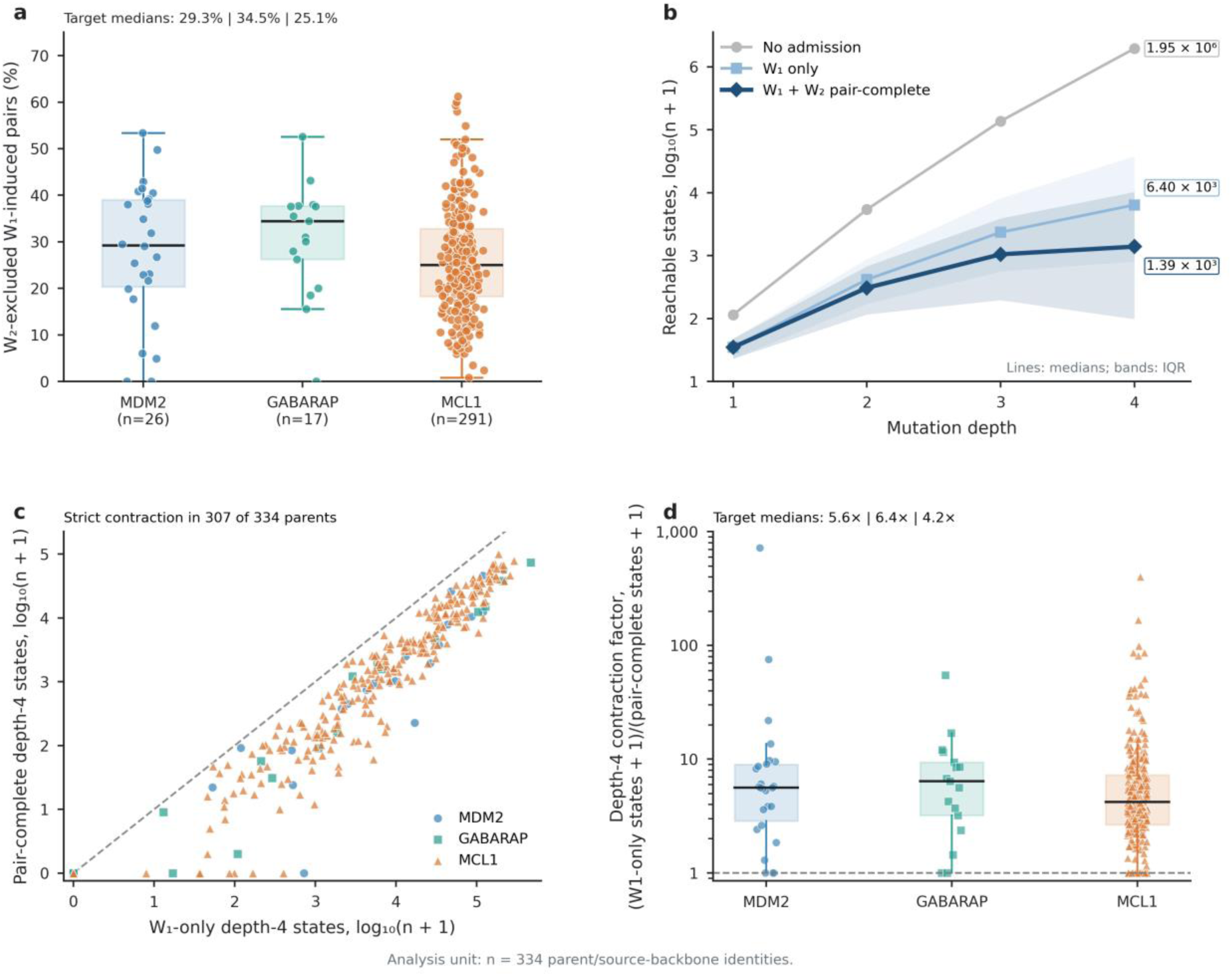
Pairwise screening excludes incompatible substitution pairs and contracts higher-order reachability. **a,** For each parent, fraction of position-distinct W₁-induced action pairs excluded by W₂. Points represent individual parent/source-backbone identities (n = 334); boxes show medians and interquartile ranges. **b,** Reachable-state counts at mutation depths 1–4 under no admission (ungated), W₁-only admission, and pair-complete W₁+W₂ admission. Lines show medians; bands show interquartile ranges. The unrestricted reference enumerates sequence combinations without structural admission and does not establish their structural feasibility. **c,** Matched per-parent depth-4 state counts under W₁-only and pair-complete admission; paired points represent the same parent across the two admission policies. **d,** Per-parent depth-4 contraction factor, defined as (W₁-only states + 1)/(pair-complete states + 1); larger values indicate greater contraction. The analysis unit is the parent/source-backbone identity (n = 334).

This pairwise incompatibility contracted the reachable sequence space progressively with mutation depth (Fig. 3b). At depth 4, the unrestricted combinatorial reference contained 1,954,815 states per parent in the absence of structural admission. The median number of reachable depth-4 states per parent was 6,400 under W₁-only admission and 1,389 under pair-complete admission. Pair-complete depth-4 counts were lower than their W₁-only counterparts for 307 of 334 parents (91.9%; Fig. 3c).

Median parent-level depth-4 contraction factors—defined as (W₁-only states + 1)/(pair-complete states + 1)—were 5.6-fold for MDM2, 6.4-fold for GABARAP, and 4.2-fold for MCL1 (Fig. 3d). Thirty-six parents (10.8%) lacked a pair-complete depth-4 state, although shallower admitted states remained reachable. Pair-level evidence therefore provided a distinct admission layer that directed higher-order property search to a smaller graph while preserving lower-depth search paths.

Within this contracted graph, Pareto-MCTS retained non-dominated candidates representing alternative trade-offs across the four model-derived scores without a fixed weighted sum (Methods). Final candidates were ranked by their minimum parent-relative improvement across the four scores. Applications requiring a larger improvement in a specific property could select a different trade-off from the retained archive.

### Prioritized outputs improve all four model-predicted property scores without negative trade-offs

We next characterized the multiproperty profiles of RF-PepTune using an independent, target-balanced set of 90 prioritized outputs (30 per target). This set was separate from the matched three-arm experiment and was used for descriptive assessment. Median parent-relative changes in the four PepTune model-native scores—permeability, solubility, non-fouling, and low hemolysis—were (+0.080, +0.082, +0.069, +0.022) for MDM2, (+0.092, +0.064, +0.059, +0.013) for GABARAP, and (+0.141, +0.082, +0.096, +0.068) for MCL1 (Fig. 4a). The minimum change across the four objectives, Δmin, was defined as the smallest parent-relative change among the four scores for each output. Δmin was positive for all 90 outputs, with target medians of 0.020 for MDM2, 0.012 for GABARAP, and 0.064 for MCL1 (Fig. 4b). The 360 parent-relative score changes (90 outputs × 4 objectives) are shown on a shared scale in target-blocked heat maps ordered by Δmin (Fig. 4c). Thus, within this independent set, prioritization improved all four model-native scores simultaneously rather than trading a negative change in one objective for gains in the others. These scores are computational predictions, not experimental measurements.

**Fig. 4.**
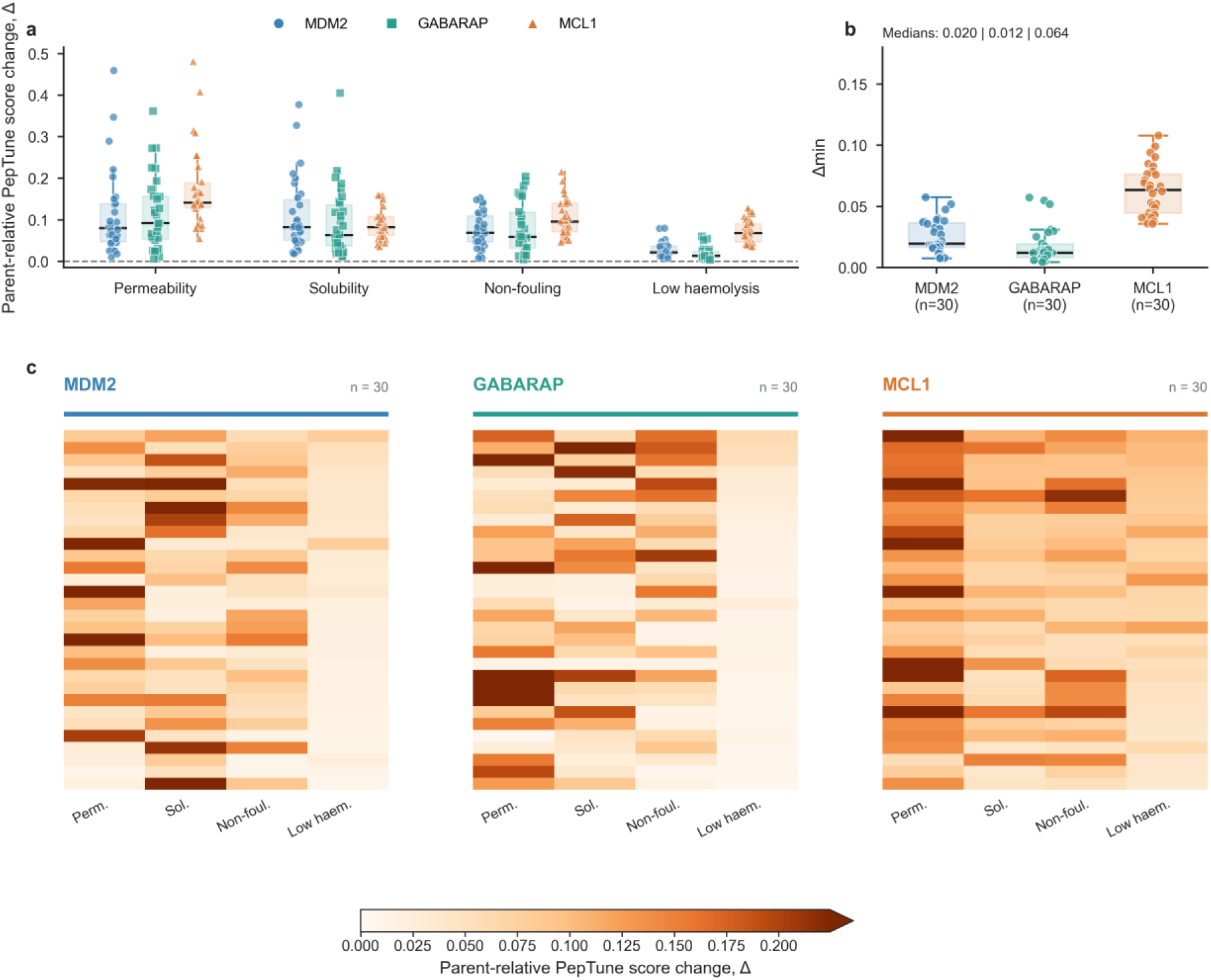
Prioritized outputs improve all four model-native property scores without negative trade-offs. **a,** Parent-relative changes in four PepTune model-native scores (permeability, solubility, non-fouling, and low hemolysis) for 30 prioritized outputs per target (MDM2, GABARAP, and MCL1). Boxes show medians and interquartile ranges; individual output points are overlaid. **b,** Minimum change across the four scores (Δmin) for each output, defined as the smallest parent-relative change among the four scores. Points represent individual outputs (n = 90; 30 per target). **c,** Target-blocked 30 × 4 heat maps sorted within each target by decreasing Δmin. Rows correspond to prioritized outputs; columns correspond to the four PepTune scores. Colors encode raw score changes on a shared linear scale; values above the pooled 95th percentile of all 360 changes use the upper extension color, with uncapped values retained in Source Data. All scores are computational predictions from PepTune, not experimental measurements.

### All prioritized endpoints pass full-sequence structural criteria under both parameterizations

To connect multiproperty score profiles to predicted structures, we selected one representative endpoint per target by deterministic ranking on Δmin, summed score change, and entry identifier (Fig. 5a–c). The selected endpoints each contained three substitutions—E3K/L12F/E15P for MDM2, S1P/E4N/L11M for GABARAP, and S2P/V16P/T17K for MCL1—with Δmin values of 0.058, 0.057, and 0.108, respectively. Their packaged complexes were rendered with explicit head-to-tail closure in the predicted target-bound geometry.

**Fig. 5.**
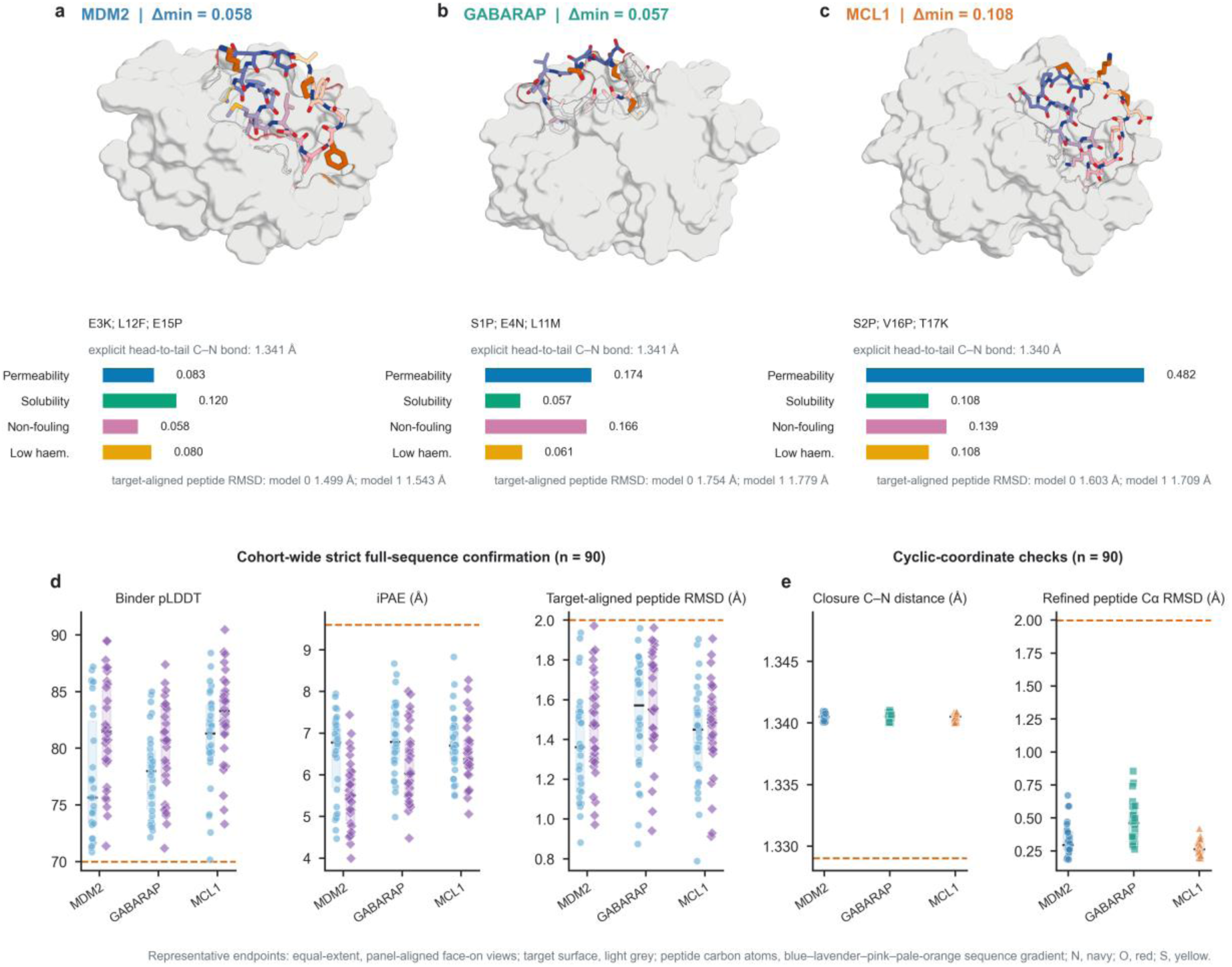
Representative cyclic-peptide endpoints and structural assessment of 90 prioritized outputs. **a–c**, Deterministically selected representative endpoint complexes for MDM2 (**a**), GABARAP (**b**), and MCL1 (**c**), rendered from packaged coordinates. Target surfaces are gray; peptide carbon atoms follow a blue–lavender–pink–pale-orange gradient, with nitrogen in navy, oxygen in red, and sulfur in yellow. Each peptide contains an explicit head-to-tail C–N bond; labeled C–N distances were recomputed from the source coordinates. Horizontal bars show parent-relative changes in the four PepTune model-native scores. **d,** Binder pLDDT, interface predicted aligned error (iPAE), and target-aligned peptide RMSD for all 90 prioritized outputs under model 0 (circles) and model 1 (diamonds). Dashed lines mark prespecified structural-assessment thresholds. **e,** Head-to-tail C–N distances and direct peptide Cα RMSDs relative to input complexes for the V2-refined inventory (n = 90; 30 per target). Dashed lines indicate comparison values. Coordinate measurements in **e** describe refined outputs, whereas **d** reports full-sequence prediction metrics from the structural-assessment workflow. Representative endpoints were selected by highest Δmin per target, then highest summed score change, then entry identifier. All constituent single substitutions and substitution pairs of the three representatives passed admission under both structural parameterizations (Supplementary Table 5). All structural metrics are computational predictions, not experimental structures.

Across all 90 prioritized outputs, binder pLDDT ranged from 70.17 to 90.46, iPAE from 4.00 to 8.83 Å, and target-aligned peptide RMSD from 0.789 to 1.972 Å under both model 0 and model 1 (Fig. 5d). Every output passed the prespecified full-sequence structural gates under both parameterizations. Because model 0 and model 1 are two parameterizations of the same structural workflow, these results represent a stringent within-workflow computational assessment rather than independent structural validation.

All 90 outputs in the harmonized V2 inventory—a coordinate-refined set used for descriptive structural quality control—passed the 11 specified computational and coordinate-audit criteria with no coordinate-parsing errors (Supplementary Fig. 1 and Supplementary Table 6). Head-to-tail C–N distances ranged from 1.33997 to 1.34103 Å, and direct peptide Cα RMSDs relative to the input complexes ranged from 0.185 to 0.854 Å (Fig. 5e). These measurements characterize closure geometry and deviation from input conformations, not accuracy against experimental structures.

## Discussion

RF-PepTune demonstrates that structural eligibility and property prioritization can be separated during local refinement of target-bound cyclic peptides. In the matched three-arm comparison, defining a pair-complete admission graph before property scoring increased joint model-defined property–structure success over single-substitution admission, while reducing property-positive endpoints that failed full-sequence structural assessment. Single-substitution admission did not determine pair admission, and pairwise restrictions propagated to higher-order sequence-space reachability. These observations support a general principle: lower-order combination evidence can define which higher-order states are eligible for property search. Because pair-complete admission changes both graph composition and graph size, the comparison estimates the net effect of the admission procedure under a fixed query budget. As a conservative rule, it may exclude higher-order sequences in which additional substitutions compensate for an inadmissible pair; full-sequence structural re-evaluation therefore remains the final assessment of each selected endpoint.

RF-PepTune builds on recent advances in target-conditioned backbone and sequence generation, cyclic-peptide structure prediction, and permeability prediction ^7–13, 20–26^. Existing methods such as AfCycDesign and RFpeptides established routes to cyclic-peptide structure prediction and target-conditioned macrocycle generation, ^7–8^ while PepTune, APCyc, and CycDiff-DPO extended generative design toward permeability and other developability objectives. ^14–16^ HFGuidedDesign and HighPlay incorporated structural prediction directly into generation or search. ^17, 19^ In contrast, RF-PepTune assigns structural and property evidence distinct algorithmic roles after a parent has been fixed: structural evidence defines the admission graph before any property score is queried, and property models then determine search priorities within that graph. This separation is not specific to cyclic peptides; it addresses a general challenge in constrained multi-objective optimization, where the feasibility of higher-order combinations cannot be inferred from lower-order components alone.

The present evidence concerns model-guided local refinement of selected parents across three targets. Property improvements were assessed using the predictors that guided optimization, and structural admission and endpoint assessment shared the same prediction workflow; predictor-specific gains and correlated structural errors therefore remain relevant to interpretation. The reported success rates measure performance under these computational criteria, and retained experimental binding and improved developability require direct measurement. Transfer to new systems also depends on parent quality and the applicability of the predictors to the designed sequences. Prospective experiments should compare matched parents and prioritized variants for target binding, permeability, solubility, hemolysis, and non-specific interactions. ^1,2, 27^ Such paired measurements would establish which predicted gains translate into improved peptide properties while retaining target binding, and would test whether structure-gated Pareto search can accelerate the development of cyclic peptide therapeutics.

## Methods

### Cyclic-peptide backbone generation and parent sequence design

Target-conditioned head-to-tail cyclic backbones were generated using the RFpeptides implementation of RFdiffusion (version [X.X]; refs. 4,8). For each target, [N] backbones were generated with the prepared receptor structure as the conditioning input, specifying a cyclic peptide of length [L] and explicit head-to-tail cyclization. Generated backbones were filtered using [criteria, e.g., binder pLDDT ≥ 70, iPAE ≤ 9.6 Å, and no steric clashes]; only qualified backbones were retained.

Fixed-backbone parent sequences were then designed with LigandMPNN (version [X.X]; refs. 5,6), which extends ProteinMPNN-style inverse folding to atomic environments. For each qualified backbone, [M] sequences were generated at temperature [T], and one parent sequence was selected by [selection criterion, e.g., highest LigandMPNN score or lowest predicted RMSD]. For each parent, the LigandMPNN-designed sequence and its matched RFpeptides backbone were used as the fixed starting point across all three comparison arms.

Receptor structures were prepared from RCSB PDB entries 1YCR (MDM2–p53 complex; 2.60 Å resolution), 7ZKR (GABARAP–Pen3-ortho stapled-peptide complex; 1.10 Å resolution), and 2PQK (MCL1–BIM BH3 complex; 2.00 Å resolution). In each case, the crystallographic partner was removed before design. The receptor structure was further processed by [removing waters, ions, and ligands; adding hydrogens; assigning protonation states; and modeling missing residues using PDBFixer or equivalent]. No additional modifications were made to the receptor during parent generation.

### Property-blind construction of W₁ and W₂ admission

For each frozen parent, a manifest specified six mutable residue indices before any PepTune score was queried. Positions were ranked by a property-blind structure-safety score:

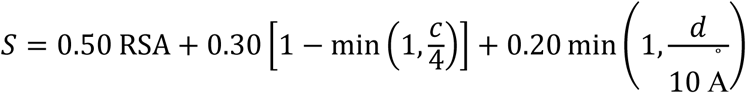

where RSA is residue solvent accessibility normalized by amino-acid-specific maximum accessibility, *c* is the number of non-adjacent intrapeptide heavy-atom contacts within 4.5 Å, and *d* is the minimum side-chain-to-target heavy-atom distance in Å. This score favors solvent-exposed positions with fewer non-adjacent intrapeptide contacts and greater side-chain separation from the target. Positions occupied by glycine or proline in the parent were deprioritized when alternative positions were available; ties were broken deterministically by sequence index.

At each selected position, the 19 non-parent canonical amino acids were enumerated, giving 114 single substitutions per parent. Each single substitution was evaluated under two parameterizations of the same structural workflow (model 0 and model 1) with three recycles. Substitutions that passed every absolute and parent-relative gate under both parameterizations defined the single-substitution admission set W₁. All position-distinct pairs induced by W₁ were then evaluated under the same two parameterizations; pairs that passed every gate defined W₂. Absolute gates were binder pLDDT ≥ 70, interface predicted aligned error (iPAE) ≤ 9.6 Å, and model-predicted target-aligned peptide RMSD ≤ 2.0 Å. Parent-relative tolerances were ΔpLDDT ≥ −5, ΔiPAE ≤ 1.0 Å, and ΔRMSD ≤ 0.5 Å.

Enumeration across the 334 parents yielded 204,186 distinct W₁-induced candidate pairs involving different residue positions (Supplementary Table 1). Each candidate pair had records under both structural parameterizations, giving 408,372 candidate-pair-by-parameterization evaluation records in the archive. PepTune scores were not used for position specification or admission-set construction.

### Pair-complete reachability and graph enumeration

For a set of substitutions *A*, a state was defined as reachable if and only if every constituent substitution belonged to W_1_ and every unordered pair of substitutions at distinct mutable positions belonged to W_2_:

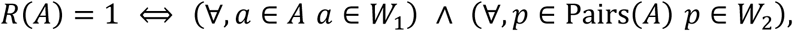

where Pairs(*A*) denotes all unordered pairs of substitutions in *A* at distinct mutable positions. A depth-3 state therefore required three admitted substitutions and all three constituent pairs; a depth-4 state required four admitted substitutions and all six constituent pairs. At mutation depth *d*, the unrestricted combinatorial reference contained 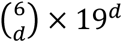 states, corresponding to the choice of *d* mutable positions and one of 19 non-parent amino acids at each position. The W₁-only count included all combinations in which each selected substitution belonged to W₁, without requiring pair-level evidence. The pair-complete count included only those combinations whose constituent substitutions all belonged to W₁ and whose induced pairs all belonged to W₂. Exact reachable-state counts were computed at depths 1–4 for all 334 frozen parents.

### PepTune property scoring and Pareto-MCTS search

Each sequence was converted to an explicit head-to-tail cyclic SMILES representation and embedded using the shared PeptideCLM/RoFormer model. Four PepTune XGBoost heads then scored permeability, solubility, non-fouling, and low hemolysis. ^15^ Search used the canonical amino-acid alphabet, a maximum Hamming distance of four, a maximum tree depth of four, and at most 256 exact PepTune queries per parent. The feasible ε-Pareto archive used ε = 0.01 and a capacity of 256. Hypervolume contributions were estimated from 512 low-discrepancy samples. Progressive-widening parameters were k = 1.5 and α = 0.5, and the exploration constant was 0.25.

Search-time property feasibility was computed from the four parent-relative score changes Δ_permeability, Δ_solubility, Δ_non-fouling, and Δ_low-hemolysis. A state was considered feasible when all four changes were at least −0.02, the larger of Δ_permeability and Δ_solubility met a target-specific primary floor, and their sum met a target-specific sum floor. Both the primary and sum floors were 0.05 for MDM2, 0.02 for GABARAP, and 0.03 for MCL1. The non-negative violation magnitude was defined as the sum of three normalized shortfalls:

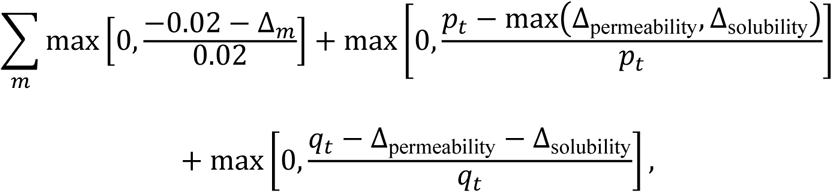

where *p_t_* and *q_t_* denote the target-specific primary and sum floors, respectively. A state was classified as feasible when this quantity was at most 1 × 10^−12^. When feasible and infeasible expansion options coexisted, a least-violating infeasible state could be explored with probability 0.15. These target-specific rules governed search-time exploration only and were distinct from the prespecified primary endpoint, whose property component required positive parent-relative changes in all four PepTune model-native scores.

### Matched three-arm comparison of structural admission policies

The matched ablation used 334 frozen parent/source-backbone identities: 26 from MDM2, 17 from GABARAP, and 291 from MCL1. For each parent, the sequence, six mutable positions, receptor and reference coordinates, amino-acid alphabet, maximum mutation depth, PepTune scorer, Pareto-MCTS implementation, hyperparameter set, 256-query cap, endpoint-selection rule, and final structural-assessment protocol were identical across arms. A frozen parent_id-to-seed map initialized an isolated Python random.Random instance for each parent–arm run. Each parent used the same seed across arms, so search trajectories differed only through the corresponding reachable graph.

All arms shared the same property-query cap, but structural prescreening introduced additional computation in the admission arms. The comparison therefore matched property-query budgets, not total computational cost.

The three arms differed only in structural admission. Ungated search allowed all 114 single substitutions and their higher-order combinations at distinct positions. W₁-only search permitted combinations of admitted single substitutions without consulting W₂. Pair-complete search required every constituent substitution to belong to W₁ and every constituent pair to belong to W₂. A neutral structural soft prior of 1.0 was applied to every reachable state. Structure-derived fields were not used for search scoring, ranking, expansion, archive quality, child selection, or tie-breaking, and the ungated and W₁-only runners did not read W₂ evidence. Runtime manifests and accessed-file records were frozen for each arm.

One endpoint of mutation depth at least two was selected per parent and arm by maximizing the minimum raw change across the four objectives, followed by maximum mean change, maximum permeability-plus-solubility change, and lexicographically maximum sequence. Graphs containing fewer than 256 evaluable states were exhausted. No endpoint satisfying the minimum mutation depth of two was reachable for one W₁-only parent and two pair-complete parents. These cases were counted as primary-endpoint failures, retaining all 334 parents in each arm’s denominator.

### Full-sequence structural re-evaluation and primary endpoint definition

Each selected endpoint was re-evaluated at the full-sequence level under two parameterizations of the same ColabDesign 1.1.1 binder-mode workflow with AlphaFold-Multimer v3 weights. Model 0 used ColabDesign model index 0 (model_1_multimer_v3; params_model_1_multimer_v3.npz), and model 1 used index 1 (model_2_multimer_v3; params_model_2_multimer_v3.npz). Both parameterizations used the binder protocol with use_multimer=True, initial_guess=True, three recycles, cyclic=True, no binder template, and the same explicit cyclic-offset procedure. Runner and configuration SHA-256 identifiers are recorded in the frozen structural-assessment protocol.

The same absolute and parent-relative gates used for W₁/W₂ construction were applied to the complete selected sequence: binder pLDDT ≥ 70, iPAE ≤ 9.6 Å, and model-predicted target-aligned peptide RMSD ≤ 2.0 Å, with parent-relative tolerances ΔpLDDT ≥ −5, ΔiPAE ≤ 1.0 Å, and ΔRMSD ≤ 0.5 Å. Parent metrics were identical across arms for all shared parents. The full-sequence structural criterion required every gate to pass under both parameterizations. Because both parameterizations belong to the same ColabDesign/AlphaFold-Multimer workflow, their joint use provides a stringent within-workflow criterion but not an independent structural validator.

The prespecified primary endpoint was joint model-defined property–structure success. Its property component required positive parent-relative changes in all four PepTune model-native scores (permeability, solubility, non-fouling, and low hemolysis). Its structural component required satisfaction of the full-sequence structural criteria under both parameterizations. Parents without a reachable endpoint were counted as primary-endpoint failures, preserving n = 334 for every arm.

### Statistical analysis of the matched three-arm comparison

The parent/source-backbone identity was the paired analysis unit (n = 334). The prespecified primary contrast was W₁+W₂ versus W₁-only admission. W₁+W₂ versus ungated search was a secondary contrast, and W₁-only versus ungated search was exploratory. Paired risk differences were reported with 100,000-replicate parent-level percentile-bootstrap 95% confidence intervals (seed 20260828). Two-sided exact McNemar tests evaluated paired binary outcomes, with Holm correction applied across the primary and secondary tests. Target-specific effects were estimated using within-target paired bootstraps. The equal-target sensitivity estimates weighted MDM2, GABARAP, and MCL1 equally. Arm-specific proportions are reported with two-sided Wilson 95% intervals.

Structural-cluster sensitivity analyses were specified post hoc, after the three-arm outcomes were known. Six complete-linkage definitions combined peptide Cα RMSD thresholds of 0.5, 1.0, or 2.0 Å with or without a contact-fingerprint criterion. Clustering was stratified by target and peptide length. For each definition, 100,000 stratified cluster-bootstrap replicates retained all parents and paired arm outcomes within each sampled cluster. Fixed original stratum weights preserved the parent-weighted estimand when combining stratum estimates. Cluster-equal estimates were reported separately as alternative estimands (Supplementary Note 2 and Supplementary Table 3).

### Frozen prioritized-output inventory and coordinate audit

The 90-structure prioritized-output inventory was generated and frozen in earlier three-target campaigns, before and independently of the matched three-arm experiment. This inventory was used for descriptive candidate characterization and post hoc coordinate-level quality control, not for the matched ablation. Candidate identities, sequences, and ranking were retained unchanged. Its source pool contained 44 MDM2, 30 GABARAP, and 151 MCL1 structures that passed the packaged-coordinate audit. Entries were ranked deterministically by the minimum parent-relative change across the four PepTune scores, then by summed change across the four scores, and finally by entry identifier; 30 structures were retained per target. The resulting 90 outputs represented 83 unique source identities because seven GABARAP outputs were additional candidates derived from sources already represented. Candidate-level property and coordinate plots therefore use n = 90 outputs and are descriptive. Cohort roles and provenance are summarized in Supplementary Table 2.

Final complexes contained an explicit head-to-tail peptide bond between the C-terminal carbon and N-terminal nitrogen and were refined with PyRosetta using the ref2015 energy function. ^32,33^ MDM2, GABARAP, and MCL1 used the same V2 refinement protocol and parameter settings (Supplementary Note 3 and Supplementary Table 4). Receptor backbone torsions and rigid-body jumps were fixed, whereas side-chain χ angles and peptide backbone torsions remained movable. Harmonic coordinate constraints were applied to peptide Cα atoms in a fixed receptor reference frame, together with explicit closure constraints. All final outputs underwent the same post hoc coordinate audit.

A post hoc uniform sensitivity analysis used the C–N distance target (1.329 Å) and estimated standard deviation (0.014 Å) recorded for the default trans-peptide TRANS link in CCP4 MON_LIB documentation, version 3.0.1 (31 May 2000); the amino-acid geometry in that library traces to Engh and Huber. ^34,35^ We defined an author-specified ±3σ sensitivity interval (1.287–1.371 Å) and applied it uniformly to every closure and sequential peptide C–N bond; all 90 frozen structures remained compliant with this interval. The ±3σ interval is a post hoc analysis tolerance, not a universal chemical acceptance interval.

## Software, reproducibility, and AI-assisted editing

Analyses were performed using Python 3.10 and 3.12, NumPy, pandas, SciPy, RDKit, PyTorch, XGBoost, OpenMM, PyRosetta, and Matplotlib. Software versions and environment specifications are recorded in the frozen protocol and accompanying code repository. The matched three-arm experiment archived a machine-readable protocol freeze, parent-to-seed map, runtime manifests, code and input hashes, per-arm accessed-file records, complete search traces, endpoint tables, structural-assessment outputs, and parent-level source data. These materials are described in the Data availability and Code availability sections and will be deposited in a public repository with a persistent identifier before publication.

OpenAI Codex was used solely for code review, figure collation, data-synchronization checks, and English-language editing. It did not generate raw data, perform statistical analyses, or determine scientific conclusions. All authors reviewed and approved the final content and take responsibility for the integrity of the work.

## Data availability

Public receptor structures are available from the RCSB Protein Data Bank under accessions 1YCR (MDM2), 7ZKR (GABARAP), and 2PQK (MCL1). Source Data for quantitative Figs. 2–5, Supplementary Fig. 1, Supplementary Tables 1–6, and Table 1 are provided as row-level CSV files; Fig. 1 is a schematic and contains no quantitative Source Data. The matched three-arm experiment evidence package contains the frozen 334-parent manifest, parent-to-seed map, per-arm search endpoints, full-sequence structural-assessment outputs, parent-level outcome table, protocol freezes, and file hashes. The prioritized-output archive contains the 90 V2-refined structures, inventory manifest, and coordinate-audit records. Supplementary audit data include the structural-clustering protocol, all six clustering definitions, cluster assignments, and representative admission records is available at https://github.com/IDLab2026/RF-peptune.

## Code availability

The code is available at https://github.com/IDLab2026/RF-peptune

## Acknowledgements

This work was supported by the Fundamental and Interdisciplinary Disciplines Breakthrough Plan of the Ministry of Education of China (Grant No. JYB2025XDXM609); National Key Research and Development Program of China (Grant No. 2022YFC2804900); National Natural Science Foundation of China (Grant Nos. 22277006, 92259302, 22677007, 22403069 and 82660813); Beijing Natural Science Foundation (Grant No. 7242195); National Talent Program (Grant Nos. CZ000935, KZ667301 and KZ0139); National Foreign Expert Program (Grant No. H20250869); National Science and Technology Major Project for Innovative Drug Research and Development (Grant No. 2026ZD1807000); 2026 Special Project for Education Development (Third Batch) - Basic Scientific Research Operating Funds for Higher Education Institutions (Grant No. CZ000713).

## Author contributions

Y.H. (Yuanchao Hou) and S.Y. (Shenwen Yang) conceived and designed the study. Y.H. (Yuanchao Hou) developed the method, implemented the code, performed the experiments, analyzed the data, and wrote the original draft. Z.L. (Zhuoyan Liu) and Y.Z. (Yixin Zhang) contributed to method development, structural screening, and data analysis. X.P. (Xinfa Peng) contributed to structural assessment and validation. Z.L. (Ziye Liu) contributed to property prediction and data analysis. M.Y. (Miyesier Yusupujiang) contributed to data curation and visualization. Y.X. (You Xu) contributed to coordinate auditing and structure refinement. J.H. (Jiaxin Hu) contributed to statistical analysis and interpretation of results. Z.L. (Zhenming Liu) and S.Y. (Shenwen Yang) supervised the project, acquired funding, and revised the manuscript. All authors discussed the results and approved the final manuscript.

## CRediT taxonomy mapping

- Conceptualization: Yuanchao Hou, Shengwen Yang
- Methodology: Yuanchao Hou, Yixin Zhang, Zhuoyan Liu, Xinfa Peng
- Software: Yuanchao Hou, Zhuoyan Liu
- Validation: Xina Peng, Ziye Liu, You Xu
- Formal analysis:Yuanchao Hou, Yixin Zhang, Jiaxin Hu
- Investigation: Yuanchao Hou, Zhuoyan Liu, Ziye Liu, Myiesier Yusupujiang
- Resources: Zhenming Liu, Shengwen Yang
- Data curation: Yixin Zhang, Myiesier Yusupujiang, You Xu
- Writing – original draft: Yuanchao Hou, Yixin Zhang
- Writing – review & editing: All authors
- Visualization: Yuanchao Hou, Myiesier Yusupujiang
- Supervision: Zhenming Liu, Shengwen Yang
- Project administration: Shengwen Yang
- Funding acquisition: Zhenming Liu, Shengwen Yang

